# pH-Dependent Membrane Binding Specificity of Synaptogyrins 1-3 Provides Mechanistic Insights into Synaptic Vesicle Regulation and Neurological Disease

**DOI:** 10.1101/2025.03.03.641025

**Authors:** Taner Karagöl, Alper Karagöl

## Abstract

Synaptogyrins (SYNGRs) are integral synaptic vesicle proteins that contribute to neurotransmitter release and synaptic plasticity. Alterations in vesicular pH, as observed in ageing and Alzheimer’s disease, may influence synaptogyrin function, yet the molecular mechanisms remain poorly understood. We compared synaptogyrin-1 (SYNGR1, pI 4.5) and synaptogyrin-3 (SYNGR3, pI 8.4), two structurally similar isoforms with distinct electrostatic properties. Using 50ns all-atom molecular dynamics simulations in realistic lipid bilayers at resting (pH 5.5) and active (pH 7.25) conditions, we examined how vesicular pH modulates protein conformation, membrane binding, and stability. Despite near-identical backbones (RMSD 1.27 Å), SYNGR1 and SYNGR3 displayed divergent pH-dependent dynamics. Comparative analysis revealed that SYNGR1’s resting state closely resembled the active state of SYNGR3, suggesting functional convergence during vesicle recycling. Multivariate amino acid profilling was conducted using homologous residue profiles. Consistent with epistatic potential, ClinVar-reported damaging variants in SYNGR1 clustered within regions of low structural mimicry, whereas SYNGR3 variants localized to conserved regions. These findings identify pH-dependent electrostatic modulation as a determinant of synaptogyrin behaviour and provide a framework for understanding their roles in synaptic vesicle cycling. The distinct conformational and mutational landscapes of SYNGR1 and SYNGR3 highlight potential mechanisms by which pH dysregulation in neurodegeneration may impair synaptic function.

## Introduction

Neuronal communication is a well-regulated process that requires the release of neurotransmitters from synaptic vesicles (Teleanu et al. 2022). This release is mediated by molecular interactions such as those between proteins and lipids within the neuron (Rizo 2022). Among the key proteins in the release of neurotransmitters are the synaptogyrins (SYNGRs), a family of integral membrane proteins that are known to be involved in synaptic vesicle trafficking (Janz et al. 1999). The pH changes upon neuronal activation are essential for the synaptic vesicle cycle (Gowrisankaran and Milosovic 2020; Farsi et al. 2017). Despite their importance, the specific mechanisms by which synaptogyrins contribute to or are affected by these processes remain incompletely understood (Gowrisankaran and Milosovic 2020).

Synaptogyrins have highly conserved features and fundamental roles, despite this, some studies show that their knockout does not significantly affect health (Park et al. 2024; Stevens 2012). High isoform counts with overlapping functions or compensatory epistatic mechanisms may be responsible for this (Park et al. 2024). Synaptogyrins also have essential functions for synaptic plasticity (Janz et al. 1999). In the brain of an Alzheimer’s disease patient, the binding interaction of Tau to synaptic vesicles is mediated by synaptogyrin-3 (SYNGR3) (Largo-Barrientos et al. 2021). Interestingly, Alzheimer’s disease and aging often involve pH changes (Decker et al. 2021).

Synaptogyrin-1 and synaptogyrin-3 are two isoforms that, although similar in structure, exhibit distinct isoelectric points (pI). This difference in pI suggests that they may interact differently with the lipid environments that have different pH levels. The synaptic vesicles maintain an acidic environment (∼pH 5.5) due to the action of proton pumps (V-ATPases) that actively transport protons (H□) into the vesicles (Forgac 2007; Farsi et al. 2017). This creates a proton gradient, which is essential for neurotransmitter storage, as the acidic pH facilitates the proton-coupled transport of neurotransmitters into the vesicles (Gowrisankaran and Milosovic 2020; Farsi et al. 2017). Upon neuronal activation, the fusion of synaptic vesicles with the plasma membrane releases their neurotransmitter cargo into the synaptic cleft (Gowrisankaran and Milosovic 2020). This process is accompanied by a shift in vesicle pH, thus ensuring a cycle of vesicle reuse and neurotransmitter recycling (Farsi et al. 2017). This pH change is likely to impact the binding affinity of proteins whose function is pH dependent. To investigate these interactions, we employed all-atom molecular dynamics simulations in lipid bilayers, allowing for the detailed study of protein interactions in a dynamic environment. We previously utilized molecular dynamics-guided lipidomic analysis of solubility enhancing variants (Karagöl et al 2024a, Johnsson et al. 2025). A realistic membrane model was a major limitation of previous simulations. In this study, lipid bilayers were constructed using a more realistic model based on previously established lipid maps (Binotti et al. 2021), according to the quantitative lipidomic analysis of mammalian synaptic vesicles (Takamori et al 2006).

Our previous work on mutational dynamics and evolutionary game theory has shown that residue substitutions in transmembrane proteins often follow patterns that optimize structural stability while permitting functional diversification (Karagöl et al. 2024b, Karagöl et al. 2025a, 2025b). Applying these frameworks here allows us to interpret synaptogyrin variability not only as a series of molecular events, but also as adaptive strategies shaped by evolutionary constraints. Mutational clustering of specific amino acids was inspected through a statistical analysis of homologous sequences. Accordingly, we utilized nonparametric correlation analysis and multivariate statistics to distinguish amino acid changes that result the diversification.

In this study, we combined molecular dynamics simulations with mutational profiling to investigate the pH-dependent behaviour of synaptogyrins. Previous studies have often been limited by simplified membrane models and by overlooking the broader evolutionary context of amino acid substitutions, we hereby employed phenotypical mapping to establish supporting clinical evidence and uncover a potential epistasis between variants of these proteins. Understanding these interactions will provide valuable insights into the molecular mechanisms that regulate synaptic proteins, potentially revealing new aspects of synaptic function and neurosecretion.

## Results and Discussions

### Comparative structural analysis of synaptogyrins 1 and 3

The two proteins in question exhibit remarkable structural similarity, reflected by a minimal root-<u>m</u>ean-square <u>d</u>eviation (RMSD) value of 1.273Å, indicating near-identical 3D conformations (Figure 1). The similar protein structures across sequences imply that the core structural elements and functional domains are preserved. On the other hand, notable non-homogeneity and clustering were observed in the regions of variation. Certain C-terminal residues showed chemically distinct variations, which could be indicative of diversification of membrane head interactions at the intracellular side (Figure 1).

**Figure 1.**
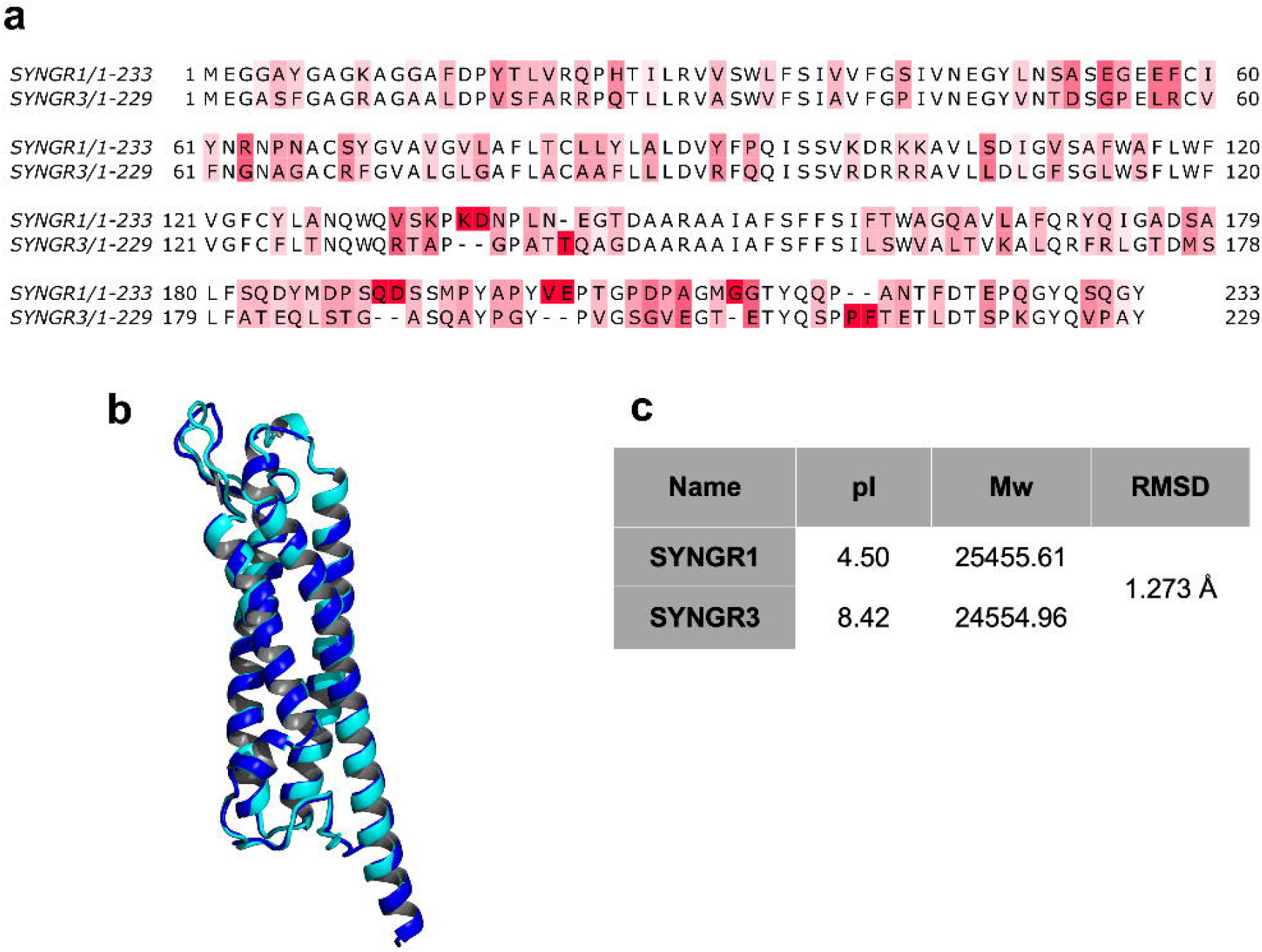
Sequence alignment and structural comparison of SYNGR1 and SYNGR3. (a) Sequence alignment of SYNGR1and SYNGR3, highlighting conserved and variable regions. Residues with high variability are coloured red, with darker red indicating greater differences. Gaps in the alignment are represented by dashes. (b) Structural superposition of SYNGR1 and SYNGR3. The SYNGR1 structure is shown in cyan and the SYNGR3 structure is in blue. (c) Summary of the isoelectric point (pI), molecular weight (Mw), and root-<u>m</u>ean-square <u>d</u>eviation (RMSD) between the two structures. SYNGR1 has a pI of 4.50 and Mw of 25455.61, while SYNGR3 has a pI of 8.42 and Mw of 24554.96, with an RMSD of 1.273 Å between the two proteins.

The high degree of structural similarity was further reinforced by significant sequence homology (Table 1). The amino acid variations in homologous sequences of the two proteins were significantly correlated (rho=0.8819, p<0.001), suggesting that they maintain a high degree of similarity in evolutionary pathways, likely due to shared structural requirements. The observed covariation could be indicative of potential compensatory mutations, where changes at one position might be balanced by alterations at another to preserve overall protein functionality. These covarying residues are likely located in structurally or functionally significant regions, such as binding sites or transmembrane domains, where precise residue interactions are critical.

**Table 1.** Residue-wise evolutionary correlations of SYNGR1 and SYNGR3.

| **Variable<sup>1</sup>** | **Spearman's rho<sup>2</sup>** | **Confidence intervals<sup>3</sup>** |
| --- | --- | --- |
| **A** | 0.8173 | 0.7428,<br>0.8795 |
| **C** | 0.6566 | 0.5020,<br>0.7695 |
| **D** | 0.7220 | 0.6247,<br>0.8120 |
| **E** | 0.6903 | 0.5891,<br>0.7794 |
| **F** | 0.8233 | 0.7455,<br>0.8860 |
| **G** | 0.8193 | 0.7488,<br>0.8761 |
| **H** | 0.5633 | 0.4251,<br>0.6779 |
| **I** | 0.7093 | 0.6108,<br>0.7887 |
| **K** | 0.7089 | 0.5832,<br>0.8058 |
| **L** | 0.7161 | 0.6137,<br>0.7954 |
| **M** | 0.6006 | 0.4698,<br>0.7021 |
| **N** | 0.6251 | 0.5141,<br>0.7244 |
| **P** | 0.6057 | 0.4862,<br>0.7070 |
| **Q** | 0.7530 | 0.6639,<br>0.8242 |
| **R** | 0.6508 | 0.5290,<br>0.7572 |
| **S** | 0.7660 | 0.6892,<br>0.8378 |
| **T** | 0.7082 | 0.6205,<br>0.7895 |
| **V** | 0.7267 | 0.6375,<br>0.8001 |
| **W** | 0.6521 | 0.4473,<br>0.8105 |
| **Y** | 0.7286 | 0.6035,<br>0.8254 |
| **Residue-wise Conservation** | 0.8819 | 0.8345,<br>0.9208 |

<sup>1</sup>Percentage distribution of amino acids at each residue position in homologous sequence alignments of SYNGR1 and SYNGR3. Residue-wise conservation is ConSurf conservation grade for each residue (1=variable 9=conserved).
<sup>2</sup>Spearman's ranked correlation ( $p < 0.001$ ).
<sup>3</sup>95% confidence intervals.

### Analysis of surface charges and relative solvent accessibilities

The similar solvent accessibility profiles may imply that they interact with their surrounding environment in comparable ways, potentially engaging with similar substrates or ligands. Accordingly, a strong correlation was found in aligned relative solvent accessibilities, rho=0.7481(95% CI=0.6357-0.8364). SYNGR1 shows a slightly more hydrophobic character overall compared to SYNGR3. The <u>m</u>olecular lipophilicity <u>p</u>otential (MLP) scores for the two structurally similar proteins reveal subtle differences in surface properties. For SYNGR3, the MLP values range from a minimum of −28.01 to a maximum of 23.17, with a mean of −1.837. In contrast, SYNGR1 shows a slightly broader range, with a minimum of −31.67, a maximum of 23.98, and a mean of −1.247. These results indicate that while both proteins have comparable hydrophobic and hydrophilic surface regions, SYNGR1 is slightly less hydrophilic (mean MLP closer to zero) and exhibits a slightly higher maximum lipophilicity than SYNGR3.

Conversely, the isoelectric points (pI) of these proteins emerge as the primary distinguishing feature between them, despite their otherwise high similarity (Figure 2). Membranes are typically negatively charged due to the presence of charged phospholipids like phosphatidylserine. Lacking a realistic membrane model was a major limitation of previous neurological protein systems we generated (Karagöl et al 2024a, Johnsson et al. 2025). In this study, lipid bilayers were constructed using a more realistic model. The model system of synaptic vesicles contains cholesterol (CHOL), phosphatidylcholine (PC), phosphatidylethanolamine (PE), phosphatidylserine (PS), sphingomyelin (SM), and phosphatidylinositol (PIns), with the following molar percentages: 40 mol%, 17 mol%, 20 mol%, 6 mol%, 4 mol%, and 1 mol%, respectively (Binotti et al. 2021; Takamori et al 2006). Among them, phosphatidylcholine and phosphatidylethanolamine are zwitterionic phospholipids, meaning they have both positive and negative charges on different parts, but their net charge is neutral. Meanwhile, phosphatidylserine and phosphatidylinositol are negatively charged. Accordingly, an increase in the positively charged areas on the protein surface could enhance electrostatic interactions with the negatively charged membrane, potentially increasing the protein’s membrane-binding affinity (Dalbey 1990). The Coulombic values for the SYNGR1 surface are as follows: minimum −12.18, mean −0.86, maximum 12.17. Meanwhile, the Coulombic values for the SYNGR3 surface are minimum −11.70, mean 1.24, maximum 12.27. For SYNGR1, the negative mean potential combined with its minimum value suggests the presence of negatively charged patches that might engage favourably with positively charged lipid headgroups.

**Figure 2.**
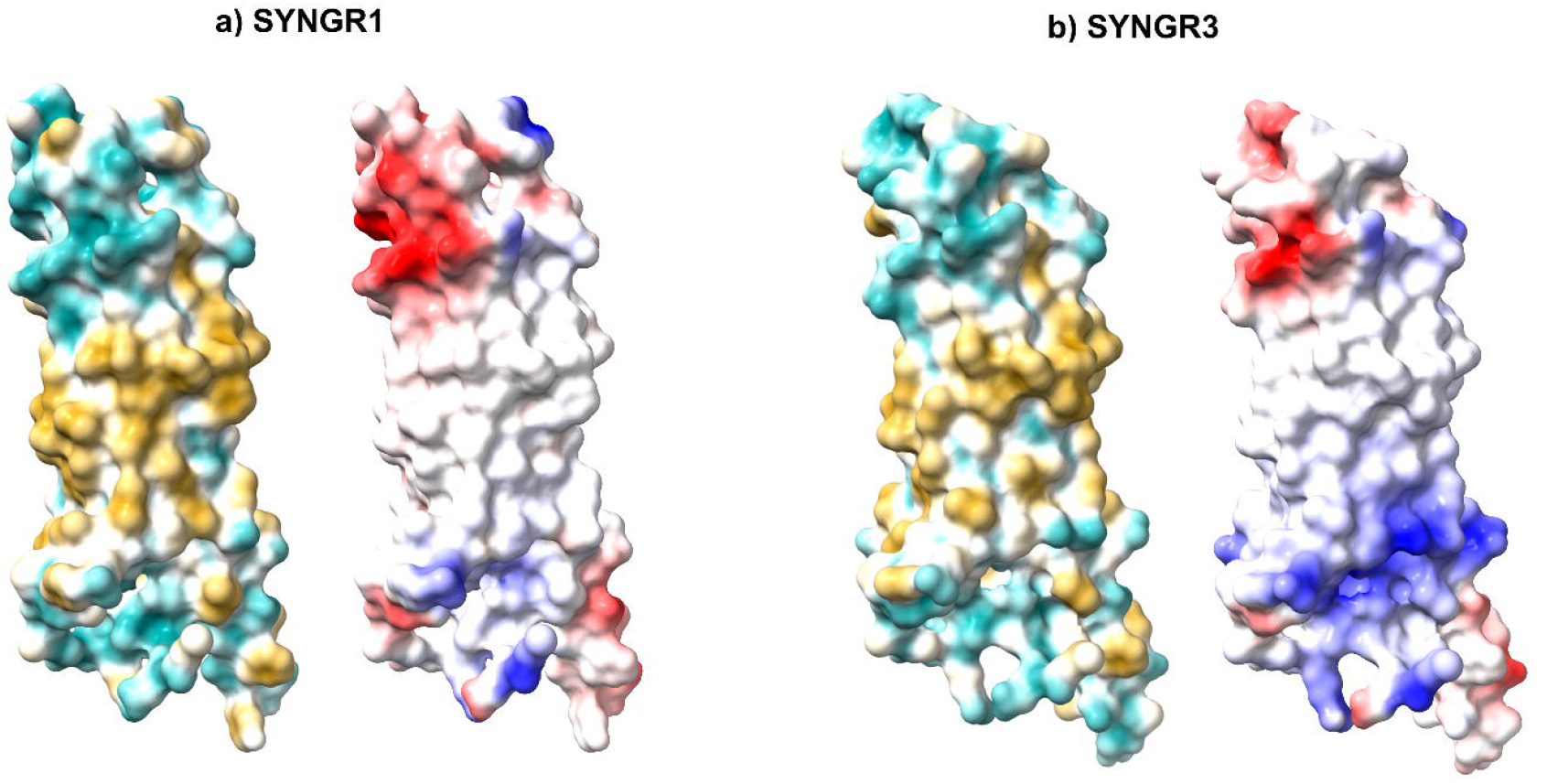
Surface profiles of SYNGR1 and SYNGR3. (a) SYNGR1 and (b) SYNGR3 are shown in molecular surface representations. The left panels depict hydrophobicity using a colour gradient (cyan indicates hydrophilic regions, and gold indicates hydrophobic regions). The right panels represent the electrostatic potential mapped onto the molecular surfaces, with red indicating negative charge, blue indicating positive charge, and white representing neutral regions. While hydrophobic surfaces show similarities, SYNGR3 has larger regions with positive charge (blue). Differences in charge distribution highlight the distinct pI values between the two isoforms.

### Dynamic behaviours of the synaptogyrins in membrane systems

The distinct isoelectric points (pI) of SYNGR1 (4.5) and SYNGR3 (8.42) provide an important context to their structural dynamics. <u>M</u>olecular <u>d</u>ynamics (MD) simulations were conducted to evaluate the structural stability of SYNGR1 under different pH conditions (Figure 3). At a physiological pH of 7.2, SYNGR1 exhibited greater conformational changes through the 50ns equilibrated trajectory, as evidenced by an average root-<u>m</u>ean-square <u>d</u>eviation (RMSD) plateau of approximately 2nm relative to its initial structure (Supplementary Figure S1). Residue-wise root-<u>m</u>ean-square fluctuations (RMSF) demonstrate a remarkable variance. This indicates that SYNGR1 undergoes a conformational change in a neutral environment. However, at a more acidic pH of 5.5, the protein exhibited extended instability, suggesting a slower transition. On the other hand, the deviations remained at 1.5nm, which is lower than the active pH of 7.5.

**Figure 3.**
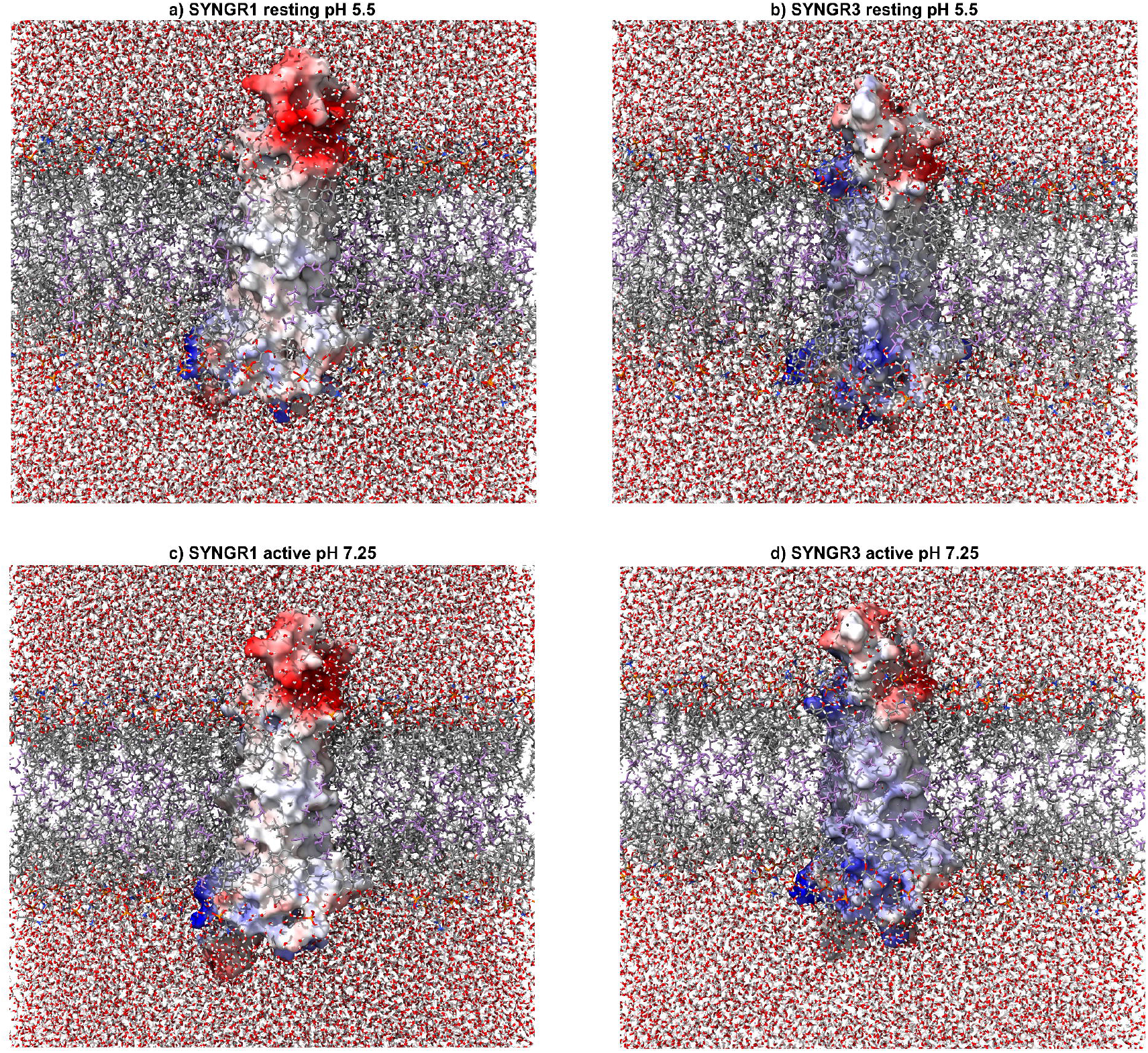
Equilibrated membrane systems of SYNGR1 (a and c) and SYNGR3 (b and d) in active and resting pH. The electrostatic potential mapped onto the molecular surfaces, with red indicating negative charge, blue indicating positive charge, and white representing neutral regions. Regions with negative charges usually located outside of the lipid bilayer. Differences in charge distribution highlight the distinct pI values between the two isoforms. Neutralizing K^+^, Cl^−^ ions (concentration=0.15M) for resting state, calcium physiologically increases in an active state and Ca^+2^ ions (concentration=0.1M) further added to active state systems (see Methods).

**Figure 4.**
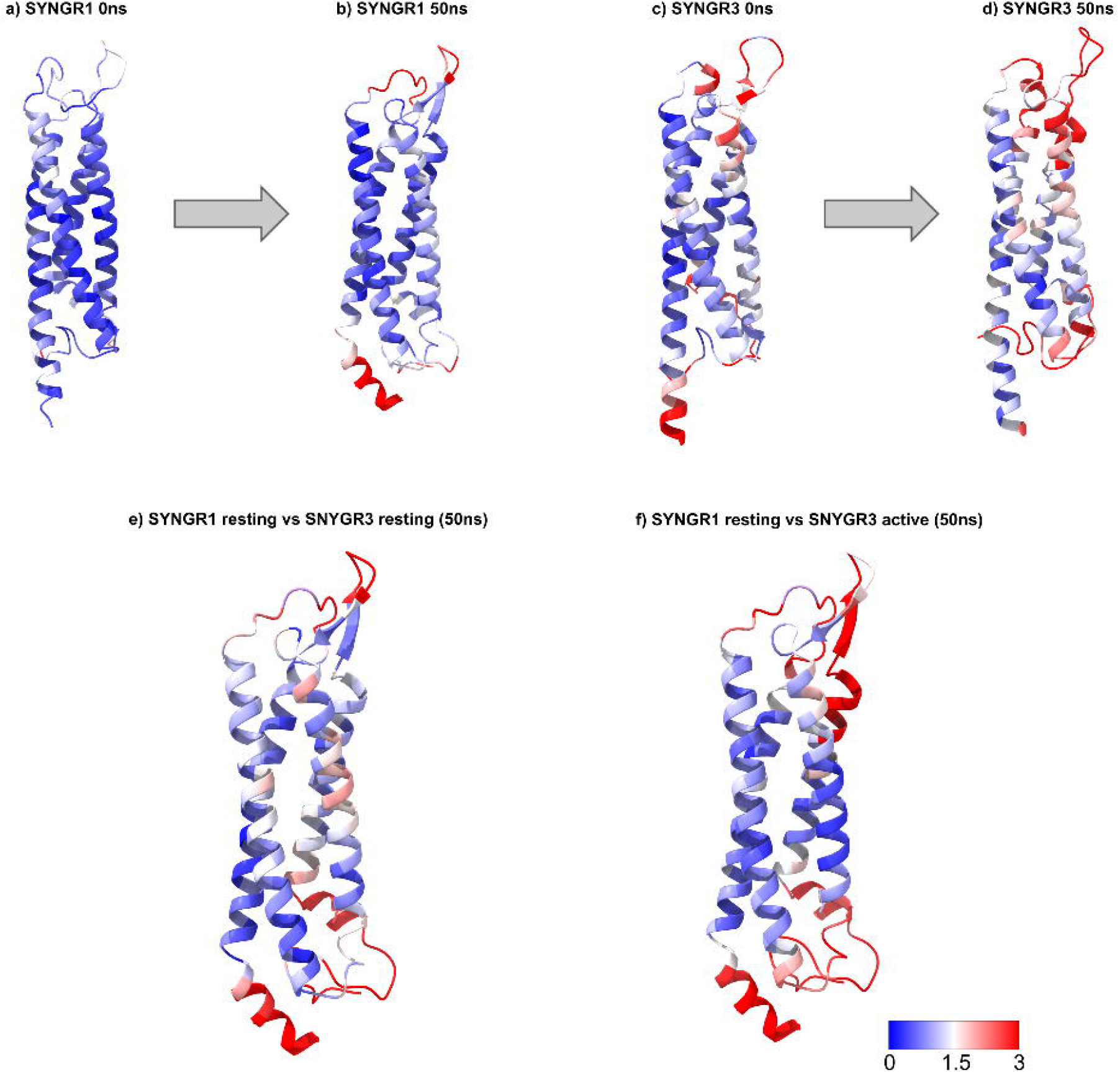
Residue-wise RMSD of SYNGR1 and SYNGR3 structures between initial and 50ns equilibrated trajectories. Panels (a) and (b) display the residue-wise deviations of SYNGR1 from its initial conformation at 0ns (a) to its equilibrated structure at 50ns (b). Similarly, panels (c) and (d) illustrate the structural progression of SYNGR3 from 0ns (c) to 50ns (d). Panels (e) and (f) compare the residue-wise RMSD between different states: (e) SYNGR1 resting pH versus SYNGR3 resting pH at 50ns, and (f) SYNGR1 resting versus SYNGR3 active pH at 50ns. The RMSD values are colour-coded, with blue indicating low deviations and red representing higher deviations, as depicted by the scale bar (0–3Å).

For SYNGR3, at pH 5.5, the protein will be positively charged overall because the pH is below the pI. This net positive charge at pH 5.5 might help the protein engage with negatively charged surfaces (such as membranes), promoting stability through electrostatic interactions. Concurrently, SYNGR3 is more stable at pH 5.5 compered to SYNGR1. At active pH (7.25), SYNGR3 reaches a plateau earlier, 20ns instead of 35ns. This could be due to better electrostatic interactions, a more favourable protonation state, or a more stable protein conformation at this pH, leading to quicker stabilization in simulations.

The observed alterations in the radius of gyration profiles between the two pH conditions further indicate that the structure of the molecule or system is sensitive to pH changes (Supplementary Figure S1). Notably, the differences in gyration axes indicate that the conformational response to pH is anisotropic, likely driven by changes in specific regions of the protein structure. As the simulation progresses, the lower pH environment (resting state) shows a smaller radius of gyration than the active state pH. The gyration axes also vary with pH changes. This is expected, as charged residues are not uniform (Figure 2, Figure 3). The distinct RMSF differences observed in residues 200–229 (C-terminus) of SYNGR3 at different pH conditions suggest a pH-dependent conformational flexibility that is not present in SYNGR1 (Supplementary Figure S2). Despite their structural similarity, SYNGR3 exhibits significant fluctuations in this region, particularly at pH 5.5, whereas C-terminal region of SYNGR1 maintains similarities across both pH conditions. At lower pH, C-terminal region in SYNGR3 might experience protonation, disrupting stabilizing intramolecular interactions and increasing local flexibility.

### Residue-wise deviations and pH dependent effects

When comparing the simulations of SYNGR1, we observe that the core transmembrane helices remain predominantly unaltered after a 50ns equilibrated simulation at active and resting pHs (RMSD below 1.5 Å). Minimal fluctuations are visible, likely due to the pI of SYNGR1 (4.5), which suggests it is negatively charged at both pH values and stabilized by interactions in its environment. Conversely, SYNGR3 undergoes more significant changes over 50ns, with red regions (RMSD over 1.5 Å), dominating the loop regions and termini, suggesting higher movement. However, at active pH (7.25), SYNGR3 reaches a plateau earlier, 20ns instead of 35ns. Some areas of the transmembrane helices also show moderate flexibility in the core region. The increased flexibility in SYNGR3 is likely due to its pI (8.42), making it less negatively charged and more adaptable to environmental changes at neutral pH compared to its behaviour at the resting pH of 5.5.

Surprisingly, the resting SYNGR1 showed more structural similarity to the active SYNGR3 than to its own resting configuration. This observation suggests that conserved structural elements critical for vesicle recycling may exist, indicating that these synaptic proteins might share key conformational features during different functional states. This similarity between resting SYNGR1 and active SYNGR3 suggests that structural elements critical for vesicle recycling (SYNGR1’s role) may be either conserved or transiently adopted by SYNGR3 during activation. This further suggests functional convergence, where active SYNGR3 mimics SYNGR1’s conformation to facilitate vesicle recycling after neurotransmitter release. Thus, our findings reinforce a more fluid, adaptable model of protein behaviour where structural elements can be transiently shared or borrowed across different protein states.

### Multivariate amino acid profiling

The divergence in pI between the two proteins could reflect differences in their amino acid compositions, particularly the presence of positively and negatively charged residues (e.g., lysine (K), arginine (R), glutamate (E), and aspartate (D)). For example, an increase in basic residues like K and R would shift the pI higher, making the protein more positively charged under physiological pH conditions, whereas an increase in acidic residues like E and D would lower the pI, resulting in a more negatively charged protein. On the other hand, the underlying variational dynamics are complex, since the correlations of homologous amino acid profiles showed that K, R, E, and D were varied, but they were not as independent as H and N (rho=0.5633 and 0.6251, respectively. If direct changes of charged residues are driving the pI divergence, we would expect them to show relatively weak correlations. A multivariate analysis was conducted using amino acids that showed lower correlations (Table 2). Interestingly, it appears that the dynamics among the amino acid varieties in the homologous sequences of SYNGR1 and SYNGR3 are asymmetrically related to H and N. For K, R, and E, removing H from SYNGR3 tends to result in the greatest decrease in correlation values, suggesting a positive interaction between H and these amino acids. On the other hand, the relationship for D appears to be enhanced by N variations in SYNGR1 homologs. D is least affected by the confounding amino acids H and N. Meanwhile, the correlation of K in homologous aligned sequences significantly decreased when controlled by H (rho=0.6655).

**Table 2.** Cofounding effects of H and N on charged amino acid (K, R, E, D) varieties in SYNGR1 and SYNGR3 homologous alignments.

| **AA** | **Control Variable<sup>1</sup>** | **Partial Spearman's rho<sup>2</sup>** | **Confidence intervals<sup>3</sup>** |
| --- | --- | --- | --- |
| **K** | H (SYNGR1) | 0.6741 | 0.5469, 0.7708 |
|  | H (SYNGR3) | 0.6655 | 0.5356, 0.7646 |
|  | N (SYNGR1) | 0.7059 | 0.5847, 0.7963 |
|  | N (SYNGR3) | 0.7071 | 0.5855, 0.7976 |
| **R** | H (SYNGR1) | 0.6358 | 0.5118, 0.7338 |
|  | H (SYNGR3) | 0.6187 | 0.4926, 0.7194 |
|  | N (SYNGR1) | 0.6503 | 0.5278, 0.7463 |
|  | N (SYNGR3) | 0.6508 | 0.5283, 0.7467 |
| **E** | H (SYNGR1) | 0.6880 | 0.5777, 0.7736 |
|  | H (SYNGR3) | 0.6773 | 0.5674, 0.7635 |
|  | N (SYNGR1) | 0.6883 | 0.5799, 0.7729 |
|  | N (SYNGR3) | 0.6913 | 0.5833, 0.7753 |
| **D** | H (SYNGR1) | 0.7206 | 0.6128, 0.8021 |
|  | H (SYNGR3) | 0.7207 | 0.6129, 0.8021 |
|  | N (SYNGR1) | 0.7026 | 0.5885, 0.7893 |
|  | N (SYNGR3) | 0.7100 | 0.5976, 0.7951 |

<sup>1</sup>Confounding variable as specific amino acids varieties in homologous sequences. For instance, N (SYNGR3) indicates N amino acid varieties in the SYNGR3 homologous sequences (aligned).
<sup>2</sup>Spearman's ranked correlation ( $p < 0.001$ ).
<sup>3</sup>95% confidence intervals.

Interestingly, among the 20 standard amino acids, histidine showed least correlation in homologous sequences (rho=0.5633). Histidine is unique among amino acids because its imidazole side chain can exist in both protonated and unprotonated states at physiological pH close to 7 (Liao et al. 2013). However, this versatility also means that histidine is more frequently substituted according to specific functional needs, depending on the protein’s environment and function (Table 1). Similarly, the amino acids M, C, N, P, R, and W have weaker correlations in homologous sequences (rho<0.66), suggesting they are less constrained and can vary more independently. The less constrained positions may provide opportunities for adaptive sequence diversification without compromising core structural integrity. In contrast, highly conserved amino acids (rho > 0.80) like A, F, G, and S are likely critical for maintaining the core structural fold or key functional interfaces.

### Clinical variant analysis and potential epistatic interactions

SYNGR1, retaining its conformation during pH differences, might be sensitive to these pathological perturbations. Accordingly, we identified that most ClinVar-reported pathological variants are located at sites that have major differences between SYNGR1 and SYNGR3 (Figure 5). SYNGR1 has more damaging variants reported compared to SYNGR3, which could be a result of differences in their evolutionary conservation or functional roles. Variants reported for Gly3, Ser36, Asn45, Ile43, Gly143, Asp145, Ala146, Arg101 are predicted to be damaging (PolyPhen score > 0.85). Most of these damaging variants are located in regions with low structural mimicry. Among them, p.Arg101Cys introduces a Cys residue capable of forming covalent disulfide bonds. The transition from a charged to a nucleophilic residue indicates an ionic interaction networks while creating the potential for aberrant cross-linking.

**Figure 5.**
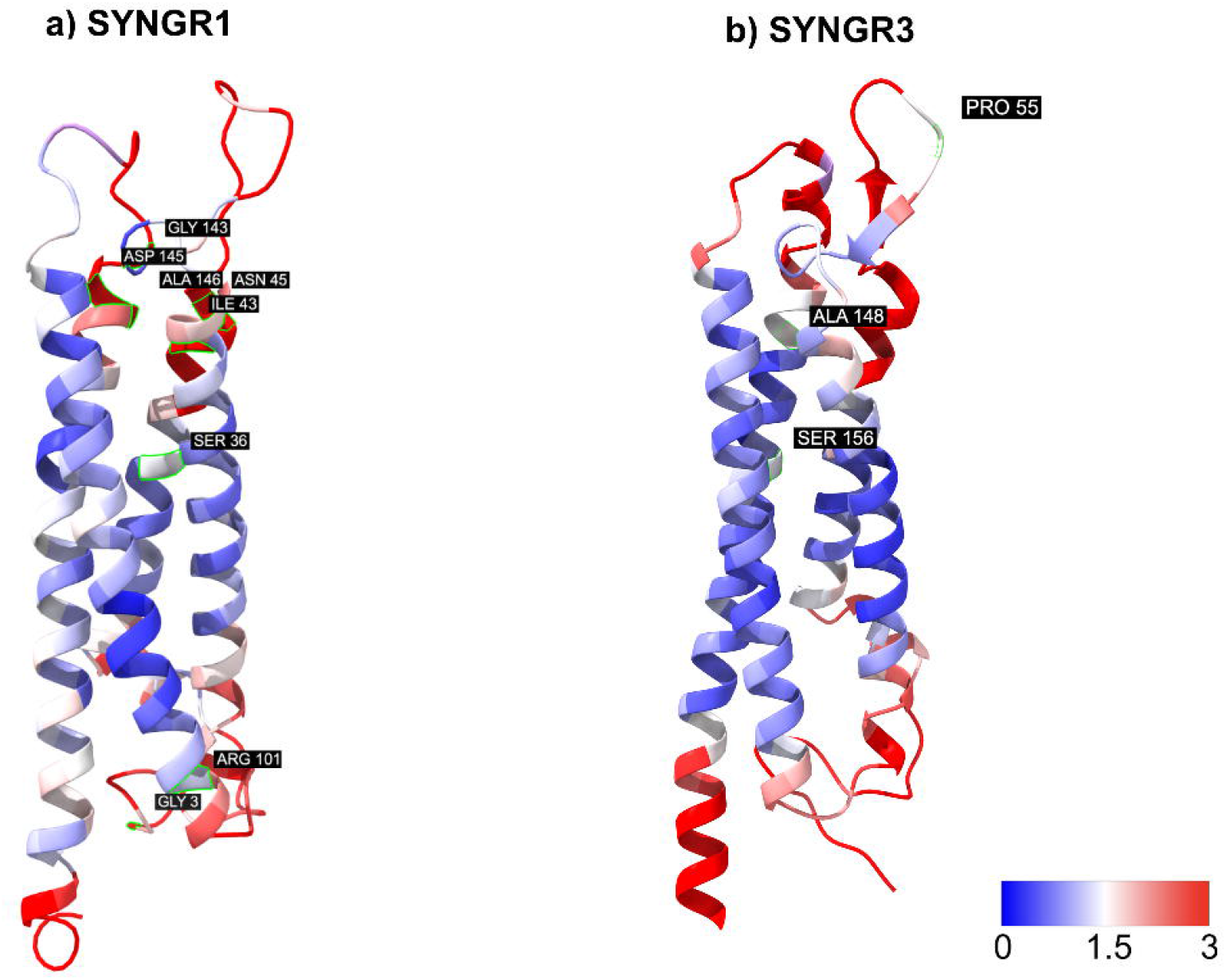
ClinVar reported variants with PolyPhen scores predicted to be damaging (0.85-1.0). Underlying colour maps compare the residue-wise RMSD between SYNGR1 resting versus SYNGR3 active pH at 50ns. The RMSD values are colour-coded, with blue indicating low deviations and red representing higher deviations, as depicted by the scale bar (0–3Å). Reported ClinVar variants for SYNGR1 were p.Gly3Arg, p.Asp145Tyr, p.Ala146Thr, p.Arg101Cys, p.Glu142Lys, p.Ser36Phe, p.Ile43Met, and p.Asn45Asp. Most of these damaging variants are located in regions with low structural mimicry. Conversely, same phenomenon could not be said for SYNGR3 variants. Reported ClinVar variants for SYNGR3 were p.Ser156Phe, p.Ala148Thr, and p.Pro55Arg.

Conversely, the same phenomenon could not be said for SYNGR3 variants. In this case, the damaging Ser156, Ala148, and Pro55 variants occur near less flexible regions, potentially affecting transmembrane stability. SYNGR3 variants are located in regions of high structural mimicry, as evidenced by RMSD maps. However, these variants demonstrate polarity changes: serine (polar, hydroxyl group) to phenylalanine (non-polar, aromatic), alanine (non-polar) to threonine (polar, hydroxyl group), and proline (non-polar, cyclic) to arginine (positively charged). This suggests that these changes may affect protein function by altering local chemical environments. Variants in SYNGR1, which predominantly affect regions of low structural mimicry, induce significant disruptions to the resting-state architecture. In contrast, SYNGR3 variants, localized to regions of high structural mimicry, potentially disrupt functional dynamics. Therefore, we suggest that active SYNGR3 adopting a more SYNGR1-like conformation could reflect a compensatory mechanism to preserve vesicle recycling while mitigating neurotransmitter release disruptions, such as in early synaptic failure in AD.

### Future scopes and the potential applications

Molecular dynamics simulations reveal that protein functionality is not solely determined by static structural characteristics but involves complex, dynamic interactions. While the 50ns simulation timeframe might be considered relatively short, it provided valuable insights into the initial stability of interactions, consistent with observations from our previous studies (Johnsson et al. 2025, Karagöl et al. 2024a, 2024c). Extending the simulations could further refine these insights. Further research could explore how pH-induced structural changes impact the functional properties of proteins, including ligand binding, allosteric regulation, or enzymatic activity. This could have direct implications for understanding diseases where pH dysregulation plays a role, such as cancer (White et al. 2017) or heart failure (Lyu et al. 2023), where the extracellular environment is altered. While this study focused on pH values of 5.5 and 7.25, extending the investigation across a broader spectrum could offer a more comprehensive understanding of the molecule’s behaviour under extreme acidic or alkaline conditions. This would be particularly relevant for systems that operate in varied biological environments, such as the gastrointestinal tract or lysosomes.

The structural resemblance between SYNGR1’s resting state and SYNGR3’s active state implies a potential evolutionary conservation of functional protein modules. These findings challenge traditional views of protein specificity and suggest a more fluid, adaptable model of protein behaviour where structural elements can be transiently shared or borrowed across different protein states. This supports the hypothesis that structural and functional modularity in proteins is an evolutionarily conserved strategy for optimizing adaptability and functionality across different environmental and cellular contexts. The observed dynamic interchangeability of structural features between SYNGR1 and SYNGR3 supports the role of evolutionary plasticity in shaping protein functionality.

## Methods

### Protein sequence alignments and related characteristics

Protein sequences for SYNGR1 and SYNGR3 were retrieved from UniProt (accession numbers O43759 and O43761) (https://www.uniprot.org) (Uniprot consortium 2023). Molecular weights, amino acid compositions, and isoelectric points were calculated using the Expasy tools (https://web.expasy.org/) (Gasteiger et al. 2005; Bjellqvist et al. 1994; Bjellqvist et al 1993).

AlphaFold3 (https://alphafoldserver.com) was used to predict structures according to default settings (Abramson 2024). The structures were then superposed using PyMOL Molecular Graphics System version 2 (https://pymol.org/2/) (Shrödinger LLC 2024). The similarities between structures were benchmarked quantitatively by calculating root-mean-square deviation (RMSD) values for all-atom RMSDs (cycles=0), ensuring the accuracy of the comparisons. Residue-wise SASA calculations and secondary structure deductions were analyzed with Stride (https://webclu.bio.wzw.tum.de/cgi-bin/stride/stridecgi.py) (Frishman and Argos 1995).

### Comparative molecular dynamics simulations

Molecular dynamics simulations were performed on the AlphaFold-predicted structures to ensure a consistent comparison. All MD simulations were performed using GROMACS 2024.3 (Abraham et al. 2015), which were executed on Google Colab, utilizing L4 GPUs, 106 GB RAM, and 43GB VRAM. To optimize computational efficiency, the simulations were parallelized across multiple cores within VM, and the source code was recompiled using all available cores with CUDA GPU for L4 GPUs according to our recent benchmarking analysis (Karagöl et al. 2024d). Configuration files and codes for the simulations are publicly available with step-by-step instructions (Karagöl et al. 2024d).

The membrane-protein systems were constructed utilizing the CHARMM-GUI web server (Wu et al. 2014; Jo et al. 2008; Jo et al. 2009), similar to our previous study on glutamate transporters (Karagöl et al. 2024a; Karagöl et al. 2024c, Johnsson et al. 2025). Spatial positions of proteins in the lipid bilayer are refined through the PPM 2.0 method, which incorporates the hydrogen-bonding profiles and anisotropic water-lipid environment (Lomize et al. 2012). Protonation states assigned based on the local pH. The lipid composition is critical for the function of synaptic vesicle proteins (Binotti et al. 2021). For our study, we used previously reviewed lipid composition profiles (Binotti et al. 2021) according to the quantitative lipidomic analysis of mammalian synaptic vesicles conducted by Takamori et al 2006. Our system of synaptic vesicles contains cholesterol (CHOL), phosphatidylcholine (PC), phosphatidylethanolamine (PE), phosphatidylserine (PS), sphingomyelin (SM), and phosphatidylinositol (PIns), with the following molar percentages: 40 mol%, 17 mol%, 20 mol%, 6 mol%, 4 mol%, and 1 mol%, respectively.

The systems were built with neutralizing K^+^, Cl^−^ ions for the resting state (concentration=0.15M), calcium levels physiologically increase in an active state and Ca^+2^ ions (concentration=0.1M) were further added to active state systems. The ions in membrane systems were determined through 2000 step Monte Carlo (MC) simulations using a primitive model. The systems were solvated in TIP3P water. CHARMM36m force field (all-atom) was used (Huang et al. 2017). The relaxation of both protein-membrane systems was conducted via multi-step minimization and equilibration. The system energies were minimized using the steepest descent method, and until the maximum forces converged to values under 1000 kJ/mol/nm. For both systems, 125-ps equilibration simulations were then performed using the standard CHARMM-GUI protocol (Wu et al. 2014; Jo et al. 2008; Jo et al. 2009). After the NVT and NPT equilibration, a 50-ns production MD simulation was run with timestamps for every 0.5ns. The Particle Mesh Ewald (PME) method was employed for electrostatics, with a cutoff distance of 1.2nm for both Coulomb and van der Waals interactions. The temperature was held at 303.15 K and pressure was held at 1 bar. The Parrinello-Rahman barostat with semi-isotropic coupling and the Nose-Hoover thermostat were utilized.

Comparative trajectory analysis was conducted for analysing the dynamical changes of both systems. The equilibrated trajectories were combined with the GROMACS gmx traj tool. The gyration radii were measured using the gmx gyrate tool. As a measure of the flexibility of residues, the residue root mean square fluctuation (RMSF) was also calculated for the Cα atoms of proteins for the full 50ns equilibrated simulation trajectory. The solvent accessible surface area (SASA) of the side chains of the protein residues was calculated according to a solvent probe radius set to 1.4Å (Eisenhaber et al. 1995). Plots that were generated through analyses were visualized utilizing Grace (https://plasma-gate.weizmann.ac.il/Grace/). Residue-wise root-<u>m</u>ean-square <u>d</u>eviation (RMSD) values were analyzed and visualized using UCSF ChimeraX version 1.8 (Meng et al. 2023).

### Analysis of Evolutionary Homology and Residue-wise Conservation Profilling

ConSurf server (https://consurf.tau.ac.il/) was utilized for generating residue-wise evolutionary conservation profiles (Yariv et al. 2023; Ashkenazy et al. 2016; Celniker et al. 2013; Landau et al. 2005). The server was run with native structures predicted by AlphaFold that were also used for RMSD calculations. The homolog search algorithm HMMER was utilized (E-value set to 0.0001). The sequences were then aligned using the MAFFT-L-INS-i method to generate a Multiple Sequence Alignment (MSA). The conservation scores were calculated using the Bayesian method, selecting the amino acid substitution model based on the best fit (default).

By comparing the percentage variations of the same amino acid residues in the aligned sequences, we statistically measured how consistently the evolutionary trends in SYNGR1 correlated with those in SYNGR3, reflecting potential co-evolutionary patterns or shared evolutionary constraints between these two proteins, in line with our previous studies (Karagöl et al. 2024b, 2025a, 2025b). The Shapiro-Wilk normality test was performed and demonstrated that the distribution of the conservation scores departed significantly from normality (Shapiro and Wilk 1965). Due to the lack of adherence to the assumptions of linearity and normality, nonparametric methods were applied, and bootstrapping was used for confidence interval calculation (Hervé 2020). Spearman’s rank correlation coefficient (Spearman’s ρ or rho) was utilized to evaluate the monotonic connection between two variables (Hervé 2020). This nonparametric method is particularly well-suited for analysing ranked data, as it does not assume a normal distribution of the variables (Kim et al. 2015), making it ideal for assessing evolutionary amino acid profiles where distributions may deviate from normality. The confounding effects of H and N on K, R, E, D frequencies were analyzed using partial Spearman coefficients. Statistical calculations were performed and visualized using R (https://www.r-project.org/) version 4.3.1 (R Core Team 2016)

### Variant analysis

Variants for *SYNGR1* and *SYNGR3* genes were obtained from large-scale sequencing projects in the Genome Aggregation Database (gnomAD v4.1.0, http://gnomad.broadinstitute.org/) (Chen et al. 2024). The Ensembl canonical transcripts for SYNGR1 and SYNGR3 were ENST00000328933.10 and ENST00000248121.7, respectively. Disease-related variants were obtained from the ClinVar database (Landrum et al. 2016) (https://www.ncbi.nlm.nih.gov/clinvar/) and *in silico* assessments of variant impact obtained via PolyPhen (http://genetics.bwh.harvard.edu/pph2/), which evaluates the likelihood of a variant affecting protein structure or function based on sequence, phylogenetic, and structural information (Adzhubei et al. 2010). PolyPhen score > 0.85 is considered likely to be damaging. Sampled variants were visualized on the residue-wise RMSD mapping of SYNGR1 and SYNGR3 using UCSF ChimeraX version 1.8 (Meng et al. 2023).

## Supplementary information (SI)

Supplementary Information.pdf

Supplementary data file featuring graphical results from the molecular dynamics simulations.

## Ethics approval

Ethics approval was not required for this computational study as it did not involve animal subjects, human participants, and identifiable data.

## Consent to participate

Not applicable. This computational study did not involve human participants.

## Consent for publication

Not applicable. This computational study did not involve human participants.

## Data availability

Each statistical and computational analysis of this study, included with step-by-step instructions where possible, are publicly available to ensure repeatability. To access detailed information on the statistical analyses, input files, and detailed results, please refer to the website: https://github.com/karagol-taner/pH-dependent-membrane-binding-synaptogyrins.

## Author Contributions

A.K. and T.K. contributed equally to this study. All authors have read and agreed to the published version of the manuscript.

## References

1) Teleanu, R.I., Niculescu, A.G., Roza, E., Vladâcenco, O., Grumezescu, A.M. and Teleanu, D.M., 2022. Neurotransmitters—key factors in neurological and neurodegenerative disorders of the central nervous system. International journal of molecular sciences, 23(11), p.5954. 10.3390/ijms23115954

2) Rizo, J., 2022. Molecular mechanisms underlying neurotransmitter release. Annual Review of Biophysics, 51(1), pp.377–408. 10.1146/annurev-biophys-111821-104732

3) Janz, R., Südhof, T.C., Hammer, R.E., Unni, V., Siegelbaum, S.A. and Bolshakov, V.Y., 1999. Essential roles in synaptic plasticity for synaptogyrin I and synaptophysin I. Neuron, 24(3), pp.687–700. 10.1016/s0896-6273(00)81122-8

4) Gowrisankaran, S. and Milosevic, I., 2020. Regulation of synaptic vesicle acidification at the neuronal synapse. IUBMB life, 72(4), pp.568–576. 10.1002/iub.2235

5) Decker, Y., Németh, E., Schomburg, R., Chemla, A., Fülöp, L., Menger, M.D., Liu, Y. and Fassbender, K., 2021. Decreased pH in the aging brain and Alzheimer’s disease. Neurobiology of aging, 101, pp.40–49. 10.1016/j.neurobiolaging.2020.12.007

6) Forgac, M., 2007. Vacuolar ATPases: rotary proton pumps in physiology and pathophysiology. Nature reviews Molecular cell biology, 8(11), pp.917–929. 10.1038/s41586-024-07610-x

7) Farsi, Z., Jahn, R., & Woehler, A. (2017). Proton electrochemical gradient: driving and regulating neurotransmitter uptake. Bioessays, 39(5), 1600240. 10.1002/bies.201600240

8) Miesenböck, G., De Angelis, D.A. and Rothman, J.E., 1998. Visualizing secretion and synaptic transmission with pH-sensitive green fluorescent proteins. Nature, 394(6689), pp.192–195. 10.1038/28190

9) Ahdut□Hacohen, R., Duridanova, D., Meiri, H. and Rahamimoff, R., 2004. Hydrogen ions control synaptic vesicle ion channel activity in Torpedo electromotor neurones. The Journal of Physiology, 556(2), pp.347–352. 10.1113/jphysiol.2003.058818

10) Park, D., Fujise, K., Wu, Y., Luján, R., Del Olmo-Cabrera, S., Wesseling, J.F. and De Camilli, P., 2024. Overlapping role of synaptophysin and synaptogyrin family proteins in determining the small size of synaptic vesicles. Proceedings of the National Academy of Sciences, 121(29), p.e2409605121. 10.1073/pnas.2409605121

11) Stevens, R.J., 2012. Genetic analysis of synaptogyrin function in the synaptic vesicle cycle (Doctoral dissertation, Massachusetts Institute of Technology).

12) Largo-Barrientos, P., Apostolo, N., Creemers, E., Callaerts-Vegh, Z., Swerts, J., Davies, C., McInnes, J., Wierda, K., De Strooper, B., Spires-Jones, T. and de Wit, J., 2021. Lowering Synaptogyrin-3 expression rescues Tau-induced memory defects and synaptic loss in the presence of microglial activation. Neuron, 109(5), pp.767–777. 10.1016/j.neuron.2020.12.016

13) Binotti, B., Jahn, R. and Pérez-Lara, Á., 2021. An overview of the synaptic vesicle lipid composition. Archives of Biochemistry and Biophysics, 709, p.108966. 10.1016/j.abb.2021.108966

14) Takamori, S., Holt, M., Stenius, K., Lemke, E.A., Grønborg, M., Riedel, D., Urlaub, H., Schenck, S., Brügger, B., Ringler, P. and Müller, S.A., 2006. Molecular anatomy of a trafficking organelle. Cell, 127(4), pp.831–846. 10.1016/j.cell.2006.10.030

15) Karagöl, A., Karagöl, T., & Zhang, S. 2024a. Molecular dynamic simulations reveal that watersoluble QTY-Variants of glutamate transporters EAA1, EAA2 and EAA3 retain the conformational characteristics of native transporters. Pharmaceutical Research, 41(10), 1965–1977. 10.1007/s11095-024-03769-0

16) Johnsson, F., Karagöl, T., Karagöl, A., & Zhang, S. 2025. Structural bioinformatic study of six human olfactory receptors and their AlphaFold3 predicted water-soluble QTY variants and OR1A2 with an odorant octanoate and TAAR9 with spermidine. QRB Discovery, 6, e2. doi:10.1017/qrd.2024.18

17) Karagöl, T., Karagöl, A., & Zhang, S. 2024b. Structural bioinformatics studies of serotonin, dopamine and norepinephrine transporters and their AlphaFold2 predicted water-soluble QTY variants and uncovering the natural mutations of L-> Q, I-> T, F-> Y and Q-> L, T-> I and Y-> F. PloS one, 19(3), e0300340. 10.1371/journal.pone.0300340

18) Karagöl, A., & Karagöl, T. 2025a. Adaptation to Solvent Environment in Toll-like Receptor 5: A Comparative Evolutionary Analysis of Membrane-bound and Soluble Forms in Epinephelus coioides. bioRxiv, 2025–02. 10.1101/2025.02.28.640895

19) Karagöl T, Karagöl T, Zhang S. 2025b. Co-evolution of alpha-helical transmembrane protein residues: large-scale variant profiling and complete mutational landscape of 2277 known PDB entries representing 504 unique human protein sequences. J Mol Evo. doi: 10.1007/s00239-025-10262-8

20) Liao, S.M., Du, Q.S., Meng, J.Z., Pang, Z.W. and Huang, R.B., 2013. The multiple roles of histidine in protein interactions. Chemistry Central Journal, 7, pp.1–12. 10.1186/1752-153X-7-44

21) Dalbey, R.E., 1990. Positively charged residues are important determinants of membrane protein topology. Trends in biochemical sciences, 15(7), pp.253–257.

22) Karagöl, A., Karagöl, T., Li, M. and Zhang, S., 2024c. Inhibitory Potential of the Truncated Isoforms on Glutamate Transporter Oligomerization Identified by Computational Analysis of Gene-Centric Isoform Maps. Pharmaceutical Research, 41(11), pp.2173–2187. 10.1007/s11095-024-03786-z

23) White, K.A., Grillo-Hill, B.K. and Barber, D.L., 2017. Cancer cell behaviors mediated by dysregulated pH dynamics at a glance. Journal of cell science, 130(4), pp.663–669. 10.1242/jcs.195297

24) Lyu, Y., Thai, P., Trinh, P., Timofeyev, V., Ginsburg, K.S., Bossuyt, J., Bers, D.M., Yamoah, E.N., Chiamvimonvat, N. and Zhang, X.D., 2023. Dysregulation of intracellular pH in the failing heart. Biophysical Journal, 122(3), p.383a. 10.1016/j.bpj.2022.11.2101

25) UniProt Consortium, 2023. UniProt: the Universal Protein Knowledgebase in 2023. Nucleic Acids Research, 51(D1), D523–D531. 10.1093/nar/gkac1052

26) Gasteiger, E., Hoogland, C., Gattiker, A., Duvaud, S.E., Wilkins, M.R., Appel, R.D. and Bairoch, A., 2005. Protein identification and analysis tools on the ExPASy server (pp. 571–607). Humana press. 10.1385/1-59259-890-0:571

27) Bjellqvist, B., Basse, B., Olsen, E. and Celis, J.E., 1994. Reference points for comparisons of two□dimensional maps of proteins from different human cell types defined in a pH scale where isoelectric points correlate with polypeptide compositions. Electrophoresis, 15(1), pp.529–539. 10.1002/elps.1150150171

28) Bjellqvist, B., Hughes, G.J., Pasquali, C., Paquet, N., Ravier, F., Sanchez, J.C., Frutiger, S. and Hochstrasser, D., 1993. The focusing positions of polypeptides in immobilized pH gradients can be predicted from their amino acid sequences. Electrophoresis, 14(1), pp.1023–1031. 10.1002/elps.11501401163

29) Abramson, J., Adler, J., Dunger, J., Evans, R., Green, T., Pritzel, A., Ronneberger, O., Willmore, L., Ballard, A.J., Bambrick, J. and Bodenstein, S.W., 2024. Accurate structure prediction of biomolecular interactions with AlphaFold 3. Nature, pp.1–3.

30) Schrödinger LLC, 2024. The PyMOL Molecular Graphics System, Version 2.5.4.

31) Frishman, D. and Argos, P., 1995. Knowledge□based protein secondary structure assignment. Proteins: Structure, Function, and Bioinformatics, 23(4), pp.566–579. 10.1002/prot.340230412

32) Yariv, B., Yariv, E., Kessel, A., Masrati, G., Chorin, A.B., Martz, E., Mayrose, I., Pupko, T. and Ben□Tal, N., 2023. Using evolutionary data to make sense of macromolecules with a “face□lifted” ConSurf. Protein Science, 32(3), p.e4582. 10.1002/pro.4582

33) Ashkenazy, H., Abadi, S., Martz, E., Chay, O., Mayrose, I., Pupko, T. and Ben-Tal, N., 2016. ConSurf 2016: an improved methodology to estimate and visualize evolutionary conservation in macromolecules. Nucleic acids research, 44(W1), pp.W344–W350. 10.1093/nar/gkw408

34) Celniker, G., Nimrod, G., Ashkenazy, H., Glaser, F., Martz, E., Mayrose, I., Pupko, T. and Ben□Tal, N., 2013. ConSurf: using evolutionary data to raise testable hypotheses about protein function. Israel Journal of Chemistry, 53(3□4), pp.199–206. 10.1002/ijch.201200096

35) Landau, M., Mayrose, I., Rosenberg, Y., Glaser, F., Martz, E., Pupko, T. and Ben-Tal, N., 2005. ConSurf 2005: the projection of evolutionary conservation scores of residues on protein structures. Nucleic acids research, 33(suppl_2), pp.W299–W302. 10.1093/nar/gki370

36) Shapiro, S.S. and Wilk, M.B., 1965. An analysis of variance test for normality (complete samples). Biometrika, 52(3-4), pp.591–611. 10.1093/biomet/52.3-4.591

37) Hervé M., 2020. RVAideMemoire: testing and plotting procedures for biostatistics. R package version 0. 9-75 https://CRAN.R-project.org/package=RVAideMemoire.

38) Kim, Y., Kim, T.H. and Ergün, T., 2015. The instability of the Pearson correlation coefficient in the presence of coincidental outliers. Finance Research Letters, 13, pp.243–257. 10.1016/j.frl.2014.12.005

39) R Core Team (2016) R: A Language and Environment for Statistical Computing. Vienna, Austria. Available from: https://www.R-project.org/. Accessed 12 June 2024

40) Abraham, M.J., Murtola, T., Schulz, R., Páll, S., Smith, J.C., Hess, B. and Lindahl, E., 2015. GROMACS: High performance molecular simulations through multi-level parallelism from laptops to supercomputers. SoftwareX, 1, pp.19–25. 10.1016/j.softx.2015.06.001

41) Karagöl, T. and Karagöl, A., 2024d. Benchmarking GROMACS on Optimized Colab Processors and the Flexibility of Cloud Computing for Molecular Dynamics. bioRxiv, pp.2024–11. 10.1101/2024.11.14.623563

42) Wu, E.L., Cheng, X., Jo, S., Rui, H., Song, K. C., Dávila-Contreras, E.M., Qi, Y., Lee, J., Monje-Galvan, V., Venable, R.M., Klauda, J.B., & Im, W., 2014. CHARMM-GUI Membrane Builder toward realistic biological membrane simulations. Journal of computational chemistry, 35(27), 1997–2004. 10.1002/jcc.23702

43) Jo, S., Kim, T., Iyer, V.G. and Im, W., 2008. CHARMM□GUI: a web□based graphical user interface for CHARMM. Journal of computational chemistry, 29(11), pp.1859–1865. 10.1002/jcc.20945

44) Jo, S., Lim, J.B., Klauda, J.B. and Im, W., 2009. CHARMM-GUI Membrane Builder for mixed bilayers and its application to yeast membranes. Biophysical journal, 97(1), pp.50–58. 10.1016/j.bpj.2009.04.013

45) Lomize, M.A., Pogozheva, I.D., Joo, H., Mosberg, H.I. and Lomize, A.L., 2012. OPM database and PPM web server: resources for positioning of proteins in membranes. Nucleic acids research, 40(D1), pp.D370–D376. 10.1093/nar/gkr703

46) Huang, J., Rauscher, S., Nawrocki, G., Ran, T., Feig, M., De Groot, B.L., Grubmüller, H. and MacKerell Jr, A.D., 2017. CHARMM36m: an improved force field for folded and intrinsically disordered proteins. Nature methods, 14(1), pp.71–73. 10.1038/nmeth.4067

47) Eisenhaber, F., Lijnzaad, P., Argos, P., Sander, C. and Scharf, M., 1995. The double cubic lattice method: Efficient approaches to numerical integration of surface area and volume and to dot surface contouring of molecular assemblies. Journal of computational chemistry, 16(3), pp.273–284. 10.1002/jcc.540160303

48) Meng, E.C., Goddard, T.D., Pettersen, E.F., Couch, G.S., Pearson, Z.J., Morris, J.H. and Ferrin, T.E., 2023. UCSF ChimeraX: Tools for structure building and analysis. Protein Science, 32(11), p.e4792. 10.1002/pro.4792

49) Chen, S., Francioli, L.C., Goodrich, J.K., Collins, R.L., Kanai, M., Wang, Q., Alföldi, J., Watts, N.A., Vittal, C., Gauthier, L.D. and Poterba, T., 2024. A genomic mutational constraint map using variation in 76,156 human genomes. Nature, 625(7993), pp.92–100. 10.1038/s41586-023-06045-0

50) Landrum, M.J., Lee, J.M., Benson, M., Brown, G., Chao, C., Chitipiralla, S., Gu, B., Hart, J., Hoffman, D., Hoover, J. and Jang, W., 2016. ClinVar: public archive of interpretations of clinically relevant variants. Nucleic acids research, 44(D1), pp.D862–D868. 10.1093/nar/gkv1222

51) Adzhubei, I.A., Schmidt, S., Peshkin, L., Ramensky, V.E., Gerasimova, A., Bork, P., Kondrashov, A.S. and Sunyaev, S.R., 2010. A method and server for predicting damaging missense mutations. Nature methods, 7(4), pp.248–249. 10.1038/nmeth0410-248

