## Supplementary information for "pH-Dependent Membrane Binding Specificity of Synaptogyrins 1-3 Provides Mechanistic Insights into Synaptic Vesicle Regulation and Neurological Disease"

### pH-Dependent Membrane Binding Specificity of Synaptogyrins 1-3 with Distinct Isoelectric Points (pI) Identified by Structural Bioinformatics and Molecular Dynamics

Taner Karagöl[1,¶,*] Alper Karagöl[1,¶,*]
[1]Istanbul University Istanbul Medical Faculty, Istanbul, Turkey
[¶]These authors contribute equally.

*To whom the correspondence should be addressed.


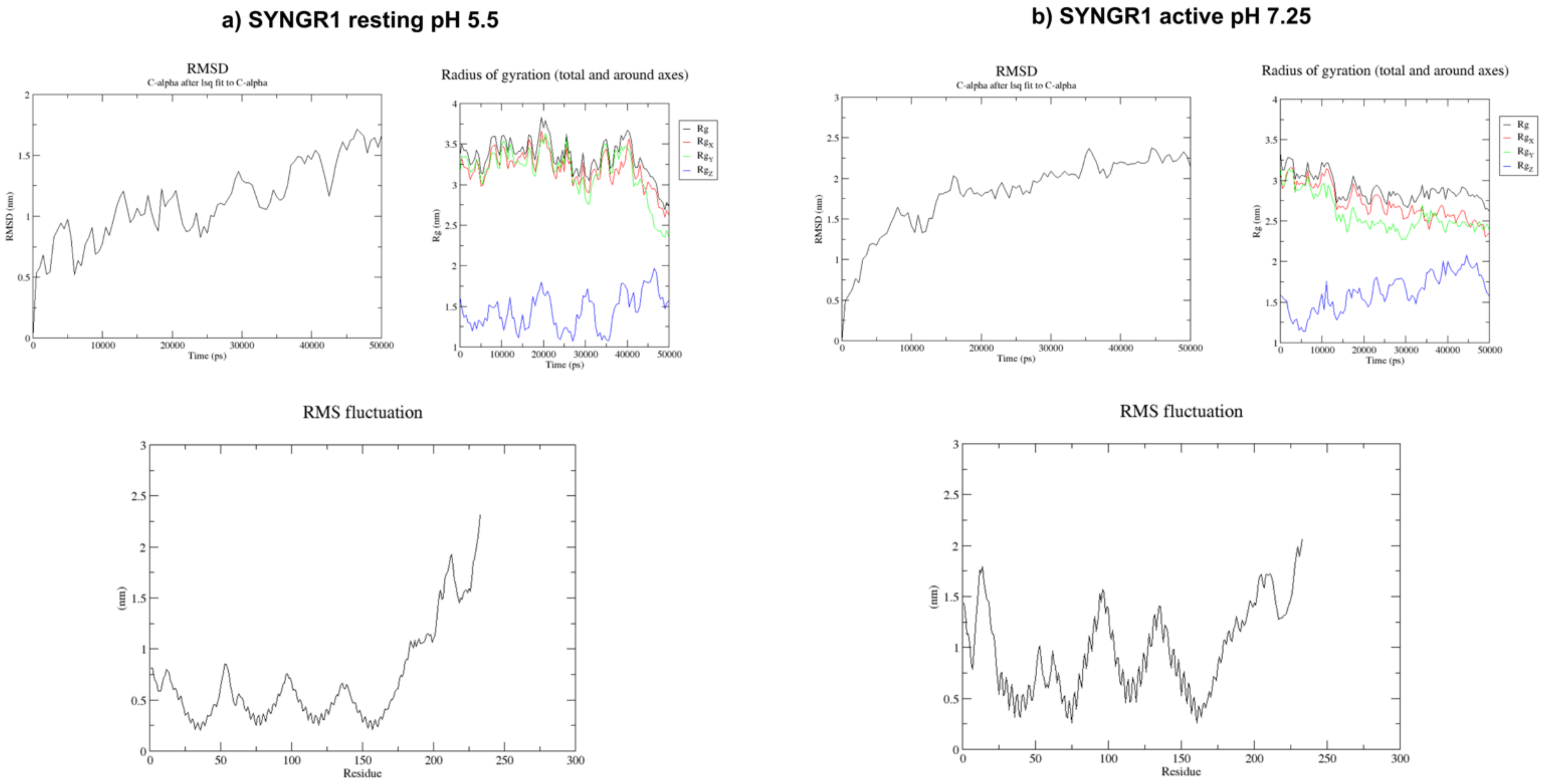


**Figure S1. Structural dynamics of SYNGR1 at different pH conditions.** (a) SYNGR1 at resting pH 5.5 and (b) at active pH 7.25. The top-left panels show the root mean square deviation (RMSD) of C-alpha atoms over 50 ns, indicating structural stability and conformational changes. The top-right panels display the radius of gyration (Rg) and its components along the X, Y, and Z axes. The bottom panels illustrate the root mean square fluctuation (RMSF) per residue. At pH 7.25, the protein exhibits higher fluctuations in specific regions, suggesting increased structural instability, whereas at pH 5.5, the protein appears more stabilized with relatively lower fluctuations.

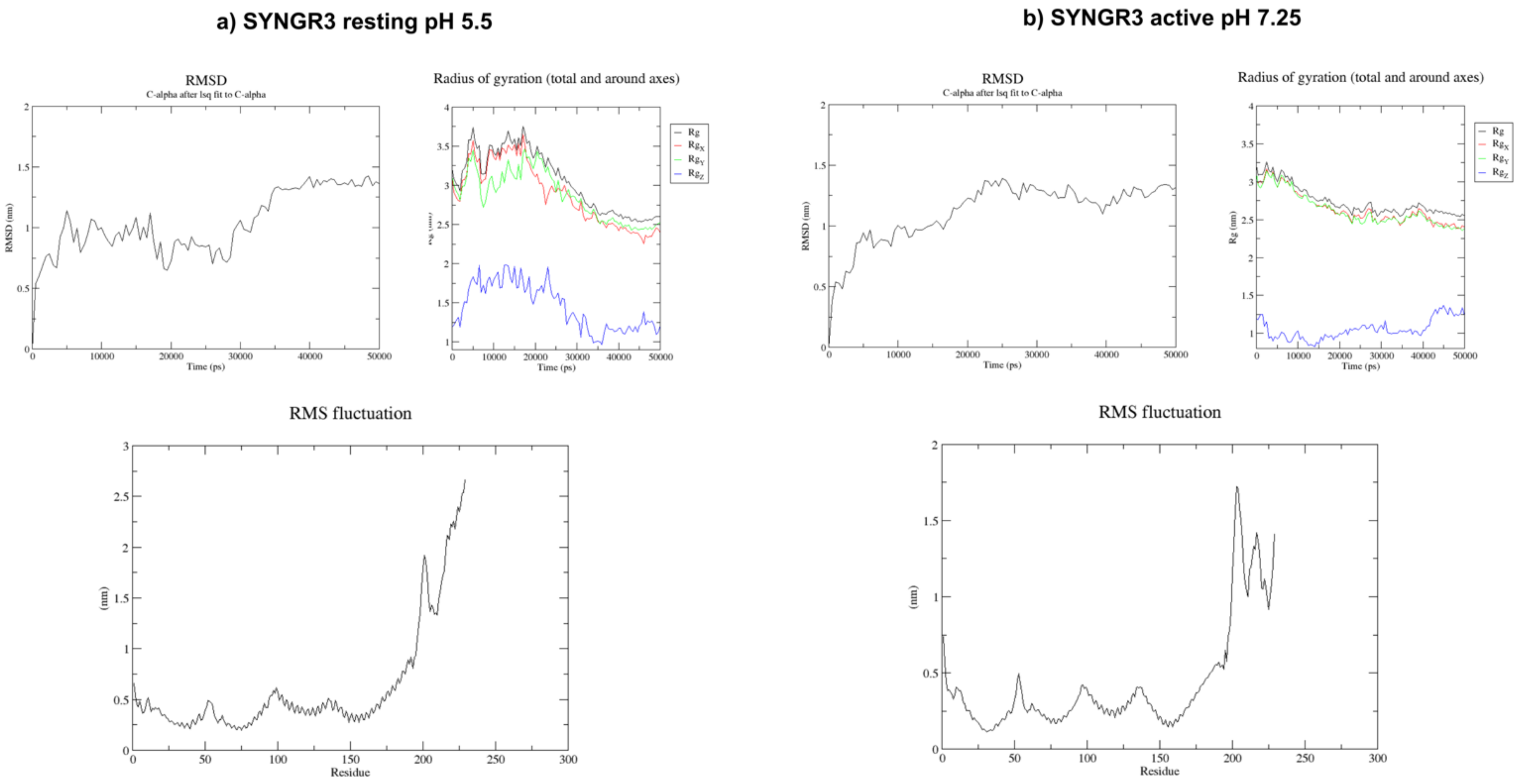


**Figure S2. Structural dynamics of SYNGR3 at different pH conditions.** (a) SYNGR3 at resting pH 5.5 and (b) at active pH 7.25. The top-left panels show the root mean square deviation (RMSD) of C-alpha atoms over 50 ns, indicating structural stability and conformational changes. The top-right panels display the radius of gyration (Rg) and its components along the X, Y, and Z axes. The bottom panels illustrate the root mean square fluctuation (RMSF) per residue. At pH 5.5, the protein exhibits higher fluctuations in specific regions, suggesting increased structural instability, whereas at pH 7.25, the protein appears more stabilized with relatively lower fluctuations.
